# Modeling vital rates and age-sex structure of Pacific arctic phocids: influence on aerial survey correction factors

**DOI:** 10.1101/2022.04.12.487942

**Authors:** Paul B. Conn, Irina S. Trukhanova

## Abstract

To estimate abundance, surveys of marine mammals often rely on samples of satellite-tagged individuals to correct counts for the proportion of animals that are unavailable to be detected. However, naïve application of this correction relies on the key assumption that availability of the tagged sample resembles that of the population. Here, we show how matrix population models can be used to estimate the stable stage structure of bearded seals (*Erignathus barbatus*), ribbon seals (*Histriophoca fasciata*), ringed seals (*Pusa hispida*), and spotted seals (*Phoca largha*) in the Bering and Chukchi Seas, and how these proportions can be used to adjust aerial survey correction factors so that they represent population-level availability. We find that correction factors ignoring age-sex composition can positively bias spotted seal abundance by an average of 13% and negatively bias ribbon seal abundance by an average of 5%. Note that we did not examine potential bias for bearded or ringed seals due to low sample sizes; as such, we urge caution in interpretation of abundance estimates for these species.

## 1. Introduction

Analysis of animal survey counts of often include corrections for availability, the proportion of animals that are detectable while surveys are being conducted (Marsh & Sinclair, 1989). For marine mammals surveyed in water, availability consists of the proportion of animals that are at the surface while the survey vessel passes (e.g. McLaren, 1961; Barlow et al., 1988); for aural surveys of songbirds, it is the proportion of birds that sing while counts are conducted (Diefenbach et al., 2007); and for aerial surveys of phocids on sea ice, it is the proportion of seals that are hauled out (Bengtson et al., 2005; Simpkins et al., 2003). In all cases, availability corrections are needed to prevent negative bias in abundance estimates (Nichols et al., 2009).

For ice-associated seals surveyed on sea ice, one possible avenue for estimating availability is to use data from satellite-linked time-depth recorder tags (TDRs) to estimate the proportion of time seals spend out of water (Bengtson et al., 2005). Indeed, availability estimates from TDRs have been used to correct survey counts of seals in the Bering (Conn et al., 2014; Ver Hoef et al. 2014) and Chukchi (Bengtson et al., 2005) seas. However, these previous efforts implicitly assumed that availability computed from the sample of tagged seals was a good approximation to that of the population – in other words, that the TDR data was not systematically biased in some way.

Recently, London et al. (2022) examined the influence of environmental variables and age-sex class on haul-out probabilities of bearded, ribbon, and spotted seals in the Bering and Chukchi seas. For ribbon and spotted seals (sample sizes for bearded seals were insufficient), they demonstrated that haul-out probabilities differed considerably by age- and sex-class. Age- and sex-specific differences in availability are logical given biological constraints, with adult females needing to spend substantial time on ice for whelping and lactation (12-18 days for bearded seals; 21-28 days for ribbon seals; 14-35 days for spotted seals; reviewed in Oftedal et al., 1987). Similarly, adult male phocids mate with females shortly after parturition, and will mate with more than one female if possible (Stirling et al., 1981). Bearded seal males also exhibit polygynous behavior and defend relatively stable territories that are advertised to females via underwater vocalization (Van Parijs and Clark, 2006); we might, then, expect adult females to spend considerably more time hauled out than adult males during pupping season, but for adults to be concentrated near sea-ice haul-out locations. By contrast, subadults of all species have no reproductive constraints, though all age classes use sea ice as a platform to undergo an annual spring molt that ranges from ≈30 days in spotted and ringed seals to ≈120 days for bearded seals (Thometz et al., 2021).

Age- and sex-mediated differences in haul-out behavior suggest a potential for bias in availability if the age-sex structure of satellite-tagged seals differs from that of the population. For ice-associated seals in the Bering and Chukchi Seas, capture operations have largely been opportunistic (Boveng et al., 2020). Adult bearded seals, for instance, are wary of humans and are particularly hard to capture on ice. Most marking operations for bearded seals have thus focused on young animals (e.g. Cameron et al., 2018; Olnes et al. 2021). Similarly, spotted seals have sometimes been caught in coastal lagoons, a process that favors capture of young individuals (e.g. Lowry et al., 1998). There is thus reason to expect that age-aggregated availability estimates for these species will be biased towards younger age classes. If younger age classes haul out less frequently than adults, naïve application of such estimates as aerial survey correction factors will likely result in positively biased abundance estimates.

In this paper, we examine the potential for bias in ice seal availability estimates when the sample of satellite-tagged seals is biased towards young individuals, and when age-sex class is ignored during estimation. We compare age aggregation to an alternative approach that attempts to estimate population-weighted availability, where age- and sex-specific availability estimates are weighted by the proportion of animals thought to be in each age-sex class. To derive these weights, we rely on stable stage distributions estimated from matrix population models (Caswell, 2001). After using data from the literature to estimate stable stage proportions for ice-associated seals, we illustrate our approach by applying both types of estimation approaches to TDR records of spotted and ribbon seals in the Bering Sea.

## 2. Methods

### 2.1 Availability models

A number of authors have used generalized linear mixed models (GLMMs; Wolfinger and O’connell, 1993) to investigate factors affecting haul-out probabilities of arctic and subarctic seals (e.g., Bengtson et al., 2005; Crawford et al., 2019; London et al., 2022; Olnes et al., 2021; Ver Hoef et al., 2010; Von Duyke et al., 2020). The GLMM framework is appealing because it allows one to include individual-level random effects, as well as autocorrelated error structures that are necessary to account for responses that are statistically dependent (as when seals haul-out consecutively for many hours). We summarized haul-out records as Bernoulli responses; for individual *i* and hour *t*, we set *Y*^*it*^=1 if individual *i* has a dry tag for ≥50% of hour *t*. According to the GLMM framework, the expected value (i.e., prediction) for individual *i* at hour *t* is a function of both fixed effects specified by the investigator (e.g., choice of explanatory predictors) and random effects which are largely governed by the specific error structure assumed and variability in the observed data. Specifically, the expected values of haul-out observations given random effects can be given as

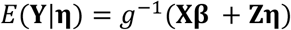

where predictive covariates are codified in a design matrix, **X** (see McCullagh & Nelder, 2019), **β** represents a vector of regression parameters, **Z** gives a design matrix for random effects that links random effects to particular observations, and *η* is a vector of random effects. The function *g*^−1^ represents an inverse link function that will typically be used to convert the linear predictor to the scale of the observations (in our application, this is the inverse logit function).

In specifying alternative models for haul-out distributions, we will focus on the fixed effects structure, **ξ** = **Xβ**, as it is here that researchers may specify different covariate effects (including sex-age effects). In particular, we consider two types of models, one with sex-age effects, and one without:

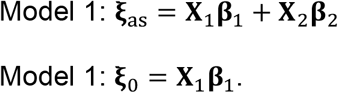

Here, **X**_2_ and **β**_2_ represent a design matrix and regression parameters for age- and sex-effects, and **X**_1_ and **β**_1_ represent a design matrix and regression coefficients for remaining covariates (e.g., weather, time-of-day), including an intercept. For specific examples of how such models can be formulated, see *Ribbon and Spotted Seal Analysis*.

The GLMM models can be fitted directly to TDR records to estimate **β**_1_ and **β**_2_ (we designate such estimates as 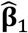 and 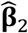). However, for aerial survey corrections we need to make predictions of the proportion of seals that were hauled out at the particular time when aerial surveys were conducted. To make predictions, let 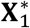 denote a design matrix where relevant entries correspond to the realized covariates at the time surveys were conducted. Availability predictions at the time of the survey can then be made using the fixed-effects structure 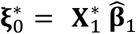 for the model without age- and sex-effects. If random effects have mean zero as is standard, a vector of availability predictions (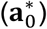) can then be generated as 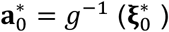 variances of predictions are slightly more complicated; see Ver Hoef et al. (2014).

However, for models with age- and sex-effects, we can only make predictions for specific age-and sex values. For instance, if we use a categorical fixed effect to represent age-sex class (e.g., young-of-year, subadult, adult female, and adult male), we can only make predictions for each of these groups separately. Without loss of generality, let us assume that we have these four age-sex classes, and denote their predictions as 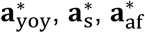, and 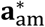, respectively. In the usual case that aerial surveys are unable to discern age-sex class (possibly they would be able to discern differences between young-of-year and other classes), it thus is necessary to average among availability predictions somehow, e.g.,

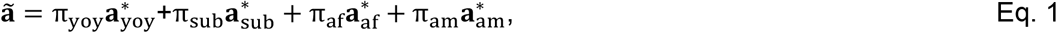

where ∑_*i*_ π_*i*_=1.0. Ideally, the weights π_*i*_ should reflect the age-sex composition of the population being surveyed. One possibility is to use stable stage proportions to help define these weights.

### 2.2 Stable stage distributions

Our strategy for computing stable stage proportions (i.e., π_yoy_, π_sub_, π_af_, π_am_) was to (1) use matrix population models (Caswell, 2001) for each species to calculation stable age proportions (ages 0-39+), and (2) to use sexual maturity schedules reported in the literature to convert stable age proportions to stable stage proportions.

In order to conduct these analyses, we first compiled data on survival, female reproduction, and sexual maturity from the literature to help calculate stable stage proportions for bearded, ribbon, ringed, and spotted seals. For survival, we used results from a hierarchical meta-analysis previously fit to a variety of phocid mortality datasets (Trukhanova et al., 2018) to produce annual survival estimates for the four species of interest. In particular, we used the same methods and data reported in Trukhanova et al. (2018), using their ribbon seal example template, to re-fit models and produce posterior predictions of survival for each of the four species. These models specify a U-shaped mortality curve (Choquet et al., 2011), with typically high mortality at the beginning of life, followed by a period of low mortality, and finally increasing mortality at older ages corresponding to senescence. We conducted analysis with JAGS 4.2.0 (Plummer, 2003), using the highest-ranked DIC model from Trukhanova et al. (2018) which included effects of subfamily, species, and dataset.

For female fecundity, we used a combination of different data sources depending on species and the amount of data available, with preference towards data collected in the Bering and Chukchi Seas (where we collect aerial survey and satellite telemetry data) and to data that were recently collected. We favored recent data since there is some indication that pregnancy rates have increased and age at sexual maturity have decreased in recent decades (Alaska Dept. of Fish & Game, 2020; Krafft et al., 2006, Quakenbush, 2020). For ringed and spotted seals, we based female reproductive and maturity on data that the Alaska Department of Fish and Game (ADFG) gathered from Alaska subsistence hunters between 2010 and 2019 (Quakenbush et al., 2020 and Alaska Dept. of Fish and Game, 2020, respectively). Reproductive assessments were limited to specimens gathered in winter and spring, when seals were reliability pregnant or had just given birth. For bearded seals, ADFG data were quite limited, so we widened the years considered to 1964-2020 and combined them with Russian data from the Bering Sea gathered in the 1980s (Fedoseev, 2000, Table 47). For ribbon seals, we based female fecundity entirely on Russian records from the Bering Sea gathered in the 1980s (Fedoseev, 2000, Table 24).

From these data, we summarized the number of specimens (*n*_*ia*_) for each species (*i*) and age class (*a*), and the number of these which were pregnant or had given birth in the year they were examined (*y*_*ia*_), as well as the number that were sexually mature (*z*_*ia*_). We truncated ages 9 and greater into a single 9+ group to reduce the likelihood of aging errors. Raw proportions by age did not necessarily increase in a smooth, monotonic fashion (particularly for ADFG data where sample sizes were small), a feature of the data that we ascribed to low sample size. In these cases, we used generalized additive models (GAMs) to smooth proportions of reproducing or mature females as a function of age. This was accomplished using the mgcv package (Wood, 2017) in the R programming environment (R Core Team 2017), using a binomial error structure:

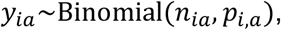

where *p*_*i,a*_, gives the proportion of females of species *i* and age class *a* that give birth in any particular year. Note that these species almost never give birth to more than a single pup per year, so *p*_*ia*_ is roughly synonymous with per capita fecundity. The GAM models incorporated smooth effects of age, and predictions from these models were incorporated into stable stage calculations (see below).

We used the notation *m*_*i,s,a*_ to denote the proportion of species *i*, sex *s*, and age *a* that are reproductively mature. For females, we used the same data sets and analysis procedures described above to estimate *m*_*i,s,a*_ from *n*_*ia*_ and *z*_*ia*_. For Russian data, sample sizes were large proportions (i.e., *m*_*i,s,a*_ = *z*_*ia*_/*n*_*ia*_). Data for the proportion of mature bearded, ribbon, and spotted seal males were taken from Tikhamorov (1966) from collections in the Bering Sea and Sea of Okhotsk. For male ringed seal maturity, we averaged proportions reported in Fedoseev (1965) and Tikhamorov (1966), weighting by sample size when they were available for both studies, and weighting each study equally when sample sizes were missing.

We used survival and reproductive values to parameterize a Leslie matrix model (Caswell, 2001). Our matrix model was structured with a post-breeding census (so that fecundity represents the product of adult survival and reproduction); it was also parameterized entirely in terms of females:

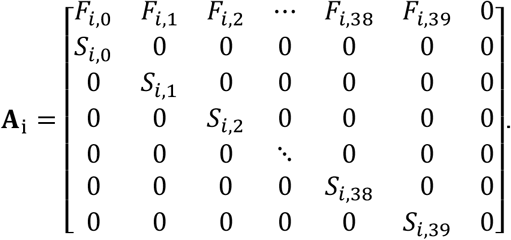

Here,*S*_*i,a*_ gives survival probability for an age *a* seal, and *F*_*i,a*_ = 0.5*p*_*i,a*_ *S*_*i,a*_ gives per capita births of female offspring for age *a* female alive at the start of the year (this assumes a sex ratio at birth of 0.5). Using this framework, we computed stable age distributions for each species as the dominant eigenvector of that species’ Leslie matrix (Caswell, 2001). All calculations were done in the R programming environment (R Core Team, 2017).

Letting *v*_*i,a*_ denote the proportion of seals of species *i* that are age *a* (determined through Leslie matrix computations), our next task is to translate these into stable stage proportions (i.e., for young-of-year, subadults, adult females, and adult males). Letting the subscript *s* = *M* denote male and *s* = *F* denote female, we have

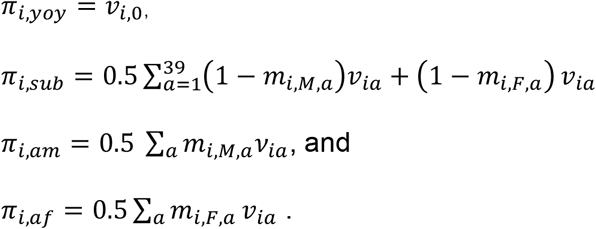

Note that these calculations implicitly assume that males and females have the same survival rates.

### 2.3 Ribbon and Spotted Seal Analysis

Having described a procedure for computing approximate stage frequencies for several Arctic phocids, we now turn our attention to analyzing possible impacts of age-sex structure on aerial survey correction factors. In particular, we reexamined haul-out behavior of 115 ribbon seals and 104 spotted seals initially analyzed by London et al. (2022). In addition to examining time-of-day and weather effects, London et al. (2022) estimated sex-age effects on haul-out probabilities, including interactions between sex-age class (young-of-year, subadult, adult female, and adult male) and day of year. In this study, we used their data and a GLMM modeling framework to (1) predict ãas in Eq. 1 using a model with an age-sex effect, and (2) predict 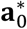 using a model without stable stage adjustments (we ran models for each species separately). To mimic aerial survey conditions during spring in the Arctic, we generated one availability prediction per day for Julian days 91-151 (April 1 -May 30 in non-leap years), at solar noon. To aid in interpretation, we omitted weather effects from GLMMs and only produced predictions for non-pups (availability of pups during Arctic spring is further complicated by the birthing process).

To see how differences in these estimates might influence abundance estimates from aerial surveys, we calculated potential bias as 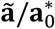. This formulation is motivated by assuming that ã is an unbiased estimate of availability, and that abundance can be reasonably calculated using a Horvitz-Thompson-like estimator (cf., Cochran 2007), i.e., 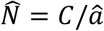 (*C* being a hypothetical aerial survey count).

### 2.4 Data and software

TDR data and haul-out predictions are currently available at https://github.com/jmlondon/berchukFits; Survival and reproductive schedules, and R code to perform stable stage estimation are currently available at https://github.com/pconn/StableStagePhocid. Both repositories will be archived to a permanent, publicly available location upon manuscript acceptance.

## 3. Results

### 3.1 Stable stage distributions

Hierarchical meta-analysis models for mortality produced estimates of annual survival for bearded, ribbon, ringed, and spotted seals that increased with age before declining due to senescence (Fig. 1a), as expected given the functional form employed (Trukhanova et al., 2018). Reproductive and sexual maturity schedules differed substantially among species (Figs. 1b-1d), with ribbon seals maturing the earliest and bearded and ringed seals maturing the slowest. These translate into very different expected stage proportions (Fig. 2). In particular, ribbon seal populations should primarily be composed of adults owing to early maturation, whereas bearded and ringed seals should have a much higher proportion of subadults.

**Figure 1.**
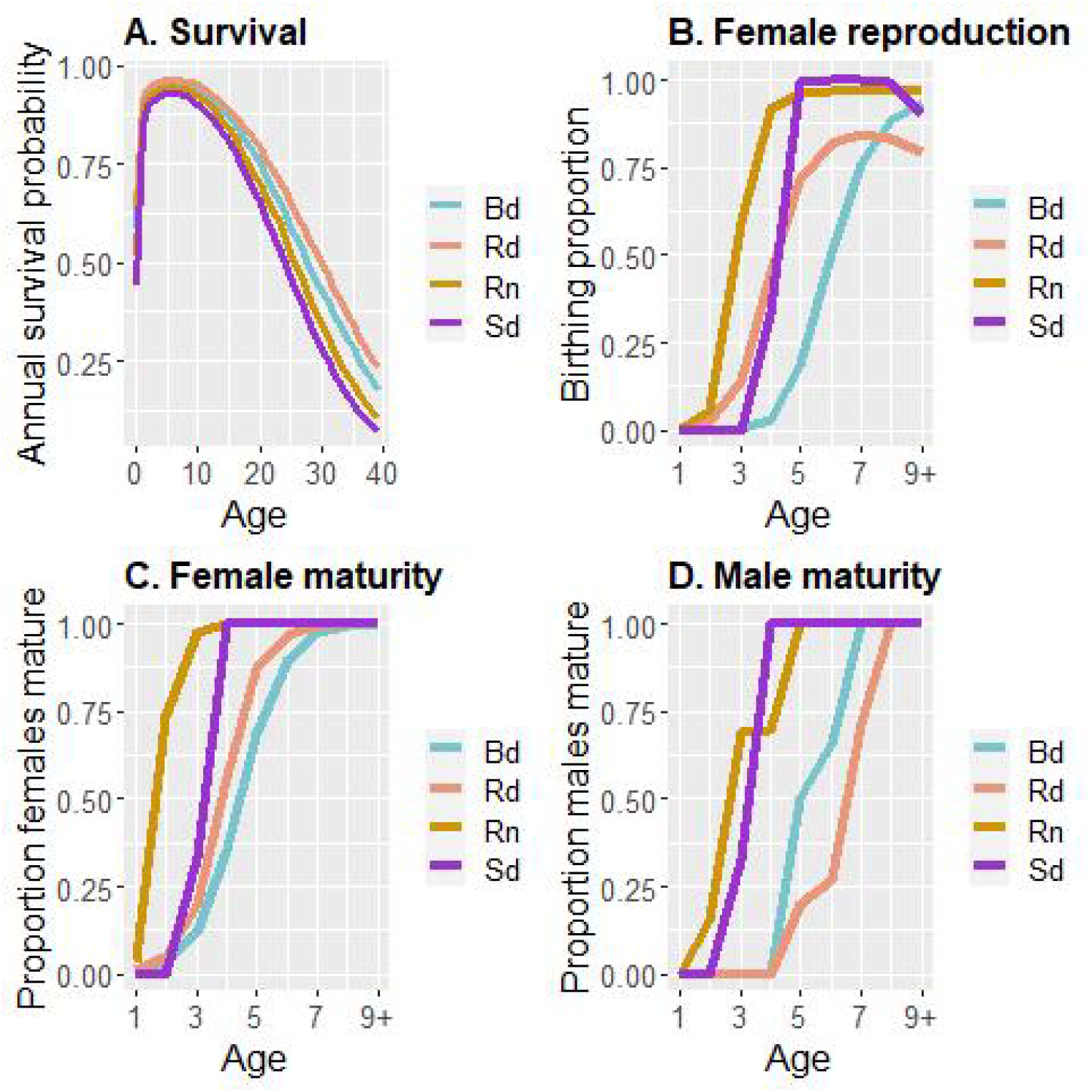
Annual survival probability, proportion of reproducing females, and proportion of sexually mature seals as a function of age for four species of ice-associated phocids (bearded seals: Bd; ringed seals: Rd; ribbon seals: Rn; and spotted seals: Sd). These estimates were used in stable stage calculations.

**Figure 2.**
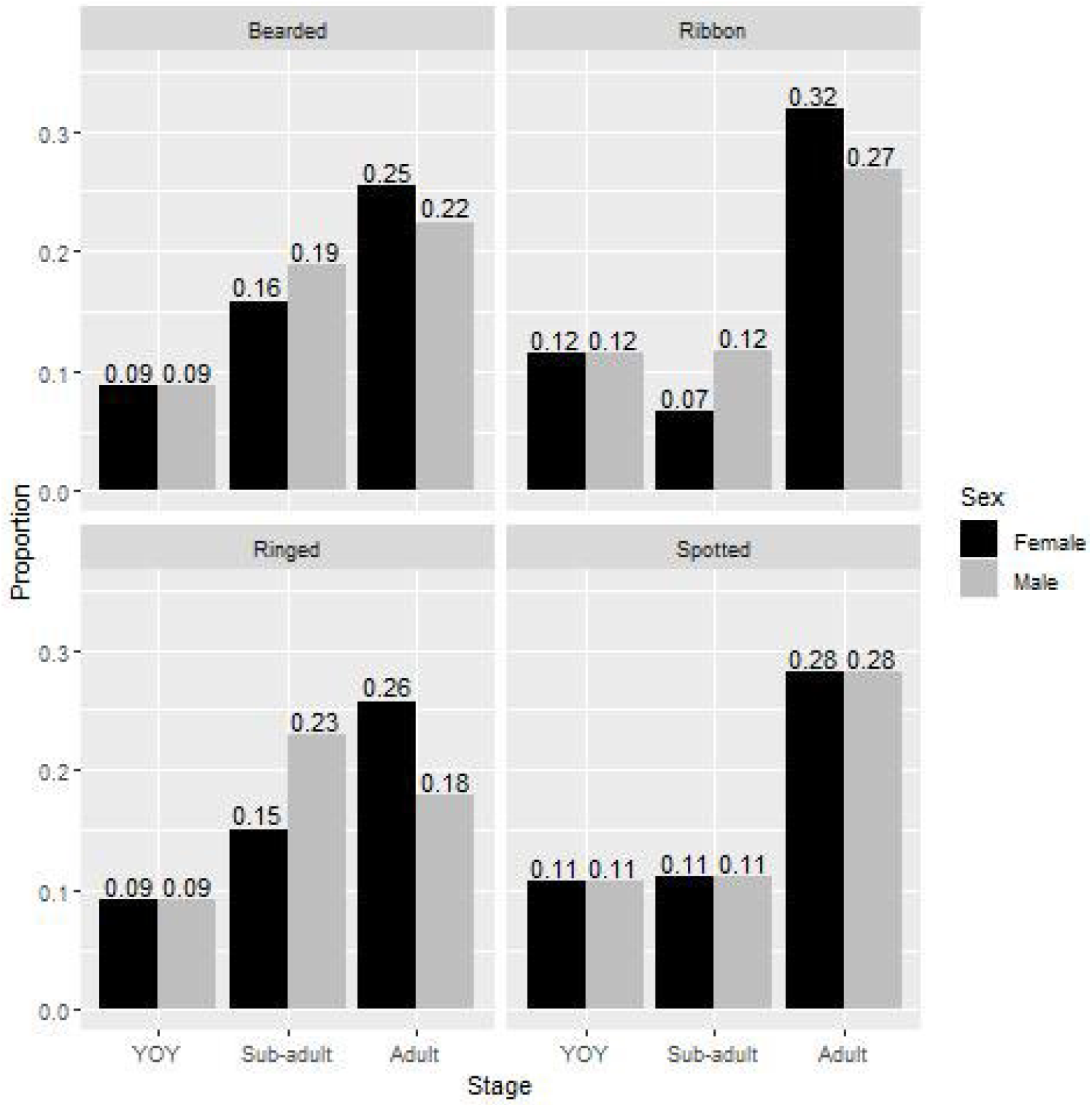
Stable stage proportions estimated from matrix population models and age-specific sexual maturity schedules for four phocid species. “YOY” stands for young-of-year.

### 3.2 Aerial survey correction factors for ribbon and spotted seals

Estimates of haul-out probabilities for spotted and ribbon seals differed depending on whether age-sex class was ignored (treating the marked sample as representative of the population) versus accounted for in predictions using stable stage weighting (Figure 3). In particular, weighted predictions were almost always higher than model predictions ignoring age-sex class. This is largely because samples of satellite tagged seals comprised higher proportions of subadults than expected for the population based on stable stage proportions. For spotted seals, 41% of non-pup telemetry observations were from subadults, whereas stable stage proportions suggested that 28% of non-pups should be subadults. For ribbon seals, 32% of non-pup records came from subadults compared to the 24% expected from stable stage proportions. Depending on the day of the survey, omitting stable-stage weighting adjustments could be expected to bias aerial survey abundance estimates of non-pups by -16 to 0% for ribbon seals (mean -5%), and by 10-20% (mean 13%) for spotted seals (Figure 3).

**Figure 3.**
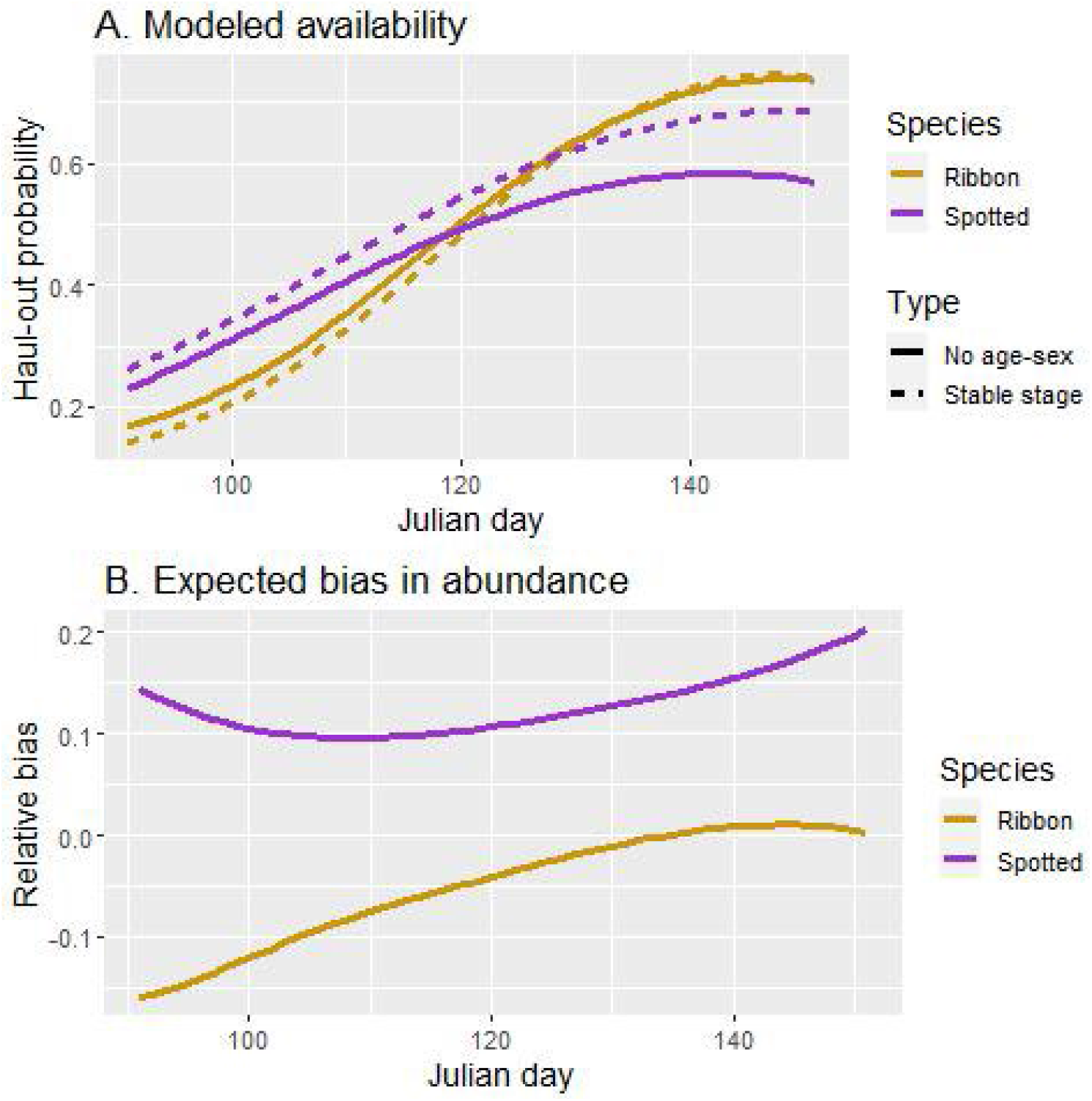
Availability and expected bias in abundance for models that do and do not adjust for age-sex effects. Panel (A) gives a comparison of predicted haul-out probabilities for subadult and adult ribbon and spotted seals for models ignoring age-sex structure (solid lines) with models that account for age-sex structure and adjust predicted haul-out behavior by stable age proportions (dashed lines). Panel (B) summarizes the expected bias in abundance estimates when age-aggregated availability estimates are used (assuming that the age- and sex-adjustment model is true).

## 4. Discussion

In this paper, we showed that estimates of availability needed for aerial survey correction factors can be biased when the age-sex composition of satellite tagged seals differs from the population and when age-aggregated availability estimates are used. We also illustrate how matrix population models and estimates of stable stage composition can be used to adjust estimates to remove these biases. Use of matrix population models and stable stage composition implicitly involves another set of assumptions – namely that life history schedules are accurate and that there have not been any major perturbations to the system (e.g., wide scale recruitment failures, extreme stochasticity in vital rates). Nevertheless, we believe stable stage proportions obtained in this manner are an improvement over the raw age proportions obtained via opportunistic sampling that might favor capture of the most vulnerable (and often younger) individuals. When assumptions underlying matrix population models are in doubt, researchers may want to conduct sensitivity analyses to investigate the effects of alternative age- and sex-class proportions on resultant availability estimates.

Where possible, our stable stage estimates relied on reproductive schedules developed from recent harvest collections (i.e., Alaska Dept. of Fish and Game, 2020; Quakenbush et al. 2020). However, ribbon seal and bearded seal reproductive schedules (and ages of male sexual maturity) relied on dated estimates from the Russian literature (i.e., Fedoseev, 1965; Fedoseev, 2000; Tikhamorov, 1966). Although these schedules work to illustrate our modeling approach, they may need to be updated for specific locations and time periods where surveys are conducted. For example, there is some indication that the age of ringed seal sexual maturity may have decreased in the last several decades (Krafft et al., 2006; Quakenbush et al., 2020); it would thus be useful to continue (and perhaps expand) sampling of Alaska Native subsistence harvests in Alaska to update our analysis with more recent data. For ringed seals, there is also some thought that mature, sexually reproducing seals preferentially select landfast ice in the Chukchi Sea, and that immature seals preferentially select pack ice habitat in the Bering Sea (Crawford et al., 2019; but see Kelly, in press). This raises an interesting question about habitat partitioning. If younger individuals select different habitats than older individuals, and if haul-out probabilities differ between the two groups, there is an argument that one should use an adult-biased correction factor in one area, and an immature-biased correction factor in the other. To make such regionally different availability predictions, researchers would need spatially explicit information on population age-sex composition, which may be difficult to collect.

Our finding that age-sex variation in seal behavior can cause biases in population dynamics parameters is not new. For instance, Härkönen et al. (1999) showed that skewed sampling of age-sex classes can result in biased estimates of population growth, survival, and fecundity in harbor seals. Similarly, Lonergan et al. (2013) illustrate how temporal variability in sex-ratios (with different haul-out probabilities for each sex) might confound inference about abundance and trend at a harbor seal haul-out site. When sample sizes allow, it thus seems prudent to account for age- and sex-based variation in behavior when applying population corrections based on biased samples of tagged seals. Our approach illustrates one possible approach for doing so. In cases where sample sizes do not allow for robust characterization of age-sex effects on availability, we advise increased caution when interpreting abundance estimates from aerial surveys of marine mammals.

## Acknowledgments

We thank our colleagues L. Quakenbush and A. Bryan (ADFG) for providing sexual maturity and reproduction data, and the Alaska Native subsistence hunters who provided the samples those data were based on. We also thank P. Boveng, M. Ferguson, J. Lee, and B. McClintock who provided comments on an earlier draft of this paper. This study was supported by NOAA, Alaska Fisheries Science Center. Any use of trade, firm, or product names is for descriptive purposes only and does not imply endorsement by the U.S Government.

## REFERENCES

Alaska Dept. of Fish and Game. (2020). Alaska Pinniped Research, Project 04: Biological monitoring of ice seals in Alaska to determine health and status of populations—diet, disease, contaminants, reproduction, body condition, growth, and age at maturity. Final Report to National Oceanographic and Atmospheric Association, Award Number: NA16NMF4390029. 180 pp + appendices.

Barlow, J., Oliver, C. W., Jackson, T. D. & Taylor, B. L. (1988). Harbor porpoise, Phocoena phocoena, abundance estimation for California, Oregon, and Washington: II. Aerial surveys. Fishery Bulletin, U.S. 86(3), 433–444.

Bengtson, J. L., Hiruki-Raring, L. M., Simpkins, M. A. & Boveng, P. L., (2005). Ringed and bearded seal densities in the eastern Chukchi Sea, 1999–2000. Polar Biology, 28(11), 833–845.

Boveng, P. L., Ziel, H. L., McClintock, B. T. & Cameron, M. F., (2020). Body condition of phocid seals during a period of rapid environmental change in the Bering Sea and Aleutian Islands, Alaska. Deep Sea Research Part II: Topical Studies in Oceanography, 181, 104904.

Cameron, M. F., Frost, K. J., Ver Hoef, J. M., Breed, G. A., Whiting, A. V., Goodwin, J. & Boveng, P. L. (2018). Habitat selection and seasonal movements of young bearded seals (Erignathus barbatus) in the Bering Sea. PloS One, 13(2), e0192743.

Caswell, H., 2001. Matrix population models (Vol. 1). Sinauer.

Choquet, R., Viallefont, A., Rouan, L., Gaanoun, K. & Gaillard, J. M. (2011). A semi-Markov model to assess reliably survival patterns from birth to death in free-ranging populations. Methods in Ecology and Evolution, 2(4), 383–389.

Cochran, W.G. (2007). Sampling techniques. John Wiley & Sons.

Conn, P. B., Ver Hoef, J. M., McClintock, B. T., Moreland, E. E., London, J. M., Cameron, M. F., Dahle, S. P. & Boveng, P.L. (2014). Estimating multispecies abundance using automated detection systems: ice-associated seals in the Bering Sea. Methods in Ecology and Evolution, 5(12), 1280–1293.

Crawford, J. A., Frost, K. J., Quakenbush, L. T. & Whiting, A. (2019). Seasonal and diel differences in dive and haul-out behavior of adult and subadult ringed seals (Pusa hispida) in the Bering and Chukchi seas. Polar Biology, 42(1), pp.65–80.

Crawford, J. A., Quakenbush, L. T. & Citta, J. J. (2015). A comparison of ringed and bearded seal diet, condition and productivity between historical (1975–1984) and recent (2003– 2012) periods in the Alaskan Bering and Chukchi seas. Progress in Oceanography, 136, 133–150.

Diefenbach, D. R., Marshall, M. R., Mattice, J. A. and Brauning, D. W. (2007). Incorporating availability for detection in estimates of bird abundance. The Auk, 124(1), 96–106.

Fedoseev, G. A. (2000). Population biology of ice-associated forms of seals and their role in the northern Pacific ecosystems. Center for Russian Environmental Policy, Russian Marine Mammal Council.

Härkönen, T., Hårding, K. C. & Lunneryd, S.G. (1999). Age-and sex-specific behaviour in harbour seals Phoca vitulina leads to biased estimates of vital population parameters. Journal of Applied Ecology, 36(5), 825–841.

Kelly B. P. (In press). The ringed seal: behavioral adaptations to seasonal ice and snow cover. In D. Costa & E. McHuron (Eds.), Ethology of earless or true seals (pp. xx), Springer.

Krafft, B. A., Kovacs, K. M., Frie, A. K., Haug, T., & Lydersen, C. (2006). Growth and population parameters of ringed seals (Pusa hispida) from Svalbard, Norway, 2002-2004. ICES Journal of Marine Science, 63, 1136–1144.

London, J. M., Conn, P. B., Hardy, S. K., Richmond, E. L., Ver Hoef, J. M., Cameron, M. F., Crawford, J. A., Von Duyke, A.L., Quakenbush, L. T. & Boveng, P. L. (2022). Haul-out behavior and aerial survey detectability of seals in the Bering and Chukchi seas. BioArxiv https://doi.org/10.1101/2022.04.07.487572.

Lonergan, M., Duck, C., Moss, S., Morris, C. & Thompson, D. (2013). Rescaling of aerial survey data with information from small numbers of telemetry tags to estimate the size of a declining harbour seal population. Aquatic Conservation: Marine and Freshwater Ecosystems, 23(1), 135–144.

Lowry, L. F., Frost, K. J., Davis, R., DeMaster, D. P. & Suydam, R. S. (1998). Movements and behavior of satellite-tagged spotted seals (Phoca largha) in the Bering and Chukchi Seas. Polar Biology, 19(4), 221–230.

Marsh, H. & Sinclair, D. F. (1989). Correcting for visibility bias in strip transect aerial surveys of aquatic fauna. The Journal of Wildlife Management, 53, 1017–1024.

McCullagh, P. & Nelder, J. A. (2019). Generalized linear models. Routledge.

McLaren, I. A., 1961. Methods of determining the numbers and availability of ringed seals in the eastern Canadian Arctic. Arctic, 14(3), 162–175.

Nichols, J. D., Thomas, L. & Conn, P. B. (2009). Inferences about landbird abundance from count data: recent advances and future directions. In D. L. Thomson, E. G. Cooch & M. J. Conroy (Eds.), Modeling demographic processes in marked populations (pp.201–235). Springer.

Oftedal, O. T., Boness, D. J. & Tedman, R. A. (1987). The behavior, physiology, and anatomy of lactation in the pinnipedia. In H. H. Genoways (Ed.), Current mammalogy (pp. 175–245). Springer.

Olnes, J., Breed, G. A., Druckenmiller, M. L., Citta, J. J., Crawford, J. A., Von Duyke, A. L. & Quakenbush, L. (2021). Juvenile bearded seal response to a decade of sea ice change in the Bering, Chukchi, and Beaufort seas. Marine Ecology Progress Series, 661, 229–242.

Quakenbush, L. T., Bryan, A., Crawford, J. & Olnes, J. (2020). Biological monitoring of ringed seals in the Bering and Chukchi seas. Final Report to National Oceanographic and Atmospheric Administration, Award Number: NA16NMF4720079. 33 pp + appendices.

R Core Team. (2017). R: A language and environment for statistical computing. R Foundation for Statistical Computing. https://www.R-project.org/.

Simpkins, M. A., Withrow, D. E., Cesarone, J. C. & Boveng, P. L. (2003). Stability in the proportion of harbor seals hauled out under locally ideal conditions. Marine Mammal Science, 19, 791–805.

Stirling, I. (1983). The evolution of mating systems in pinnipeds. In J. F. Eisenberg & D. G. Kleiman (Eds.) Recent advances in the study of mammalian behavior (pp.489–527), Special Pub. No. 7, American Society of Mammalogists.

Thometz, N. M., Hermann-Sorensen, H., Russell, B., Rosen, D. A. & Reichmuth, C. (2021). Molting strategies of Arctic seals drive annual patterns in metabolism. Conservation Physiology, 9(1), p.coaa112.

Van Parijs, S. M. & Clark, C. W. (2006). Long-term mating tactics in an aquatic-mating pinniped, the bearded seal, Erignathus barbatus. Animal Behaviour, 72(6), 1269–1277.

Ver Hoef, J. M., Cameron, M. F., Boveng, P. L., London, J. M. & Moreland, E. E. (2014). A spatial hierarchical model for abundance of three ice-associated seal species in the eastern Bering Sea. Statistical Methodology, 17, 46–66.

Ver Hoef, J. M., London, J. M. & Boveng, P. L. (2010). Fast computing of some generalized linear mixed pseudo-models with temporal autocorrelation. Computational Statistics, 25(1), 39–55.

Von Duyke, A. L., Douglas, D. C., Herreman, J. K. & Crawford, J. A. (2020). Ringed seal (Pusa hispida) seasonal movements, diving, and haul-out behavior in the Beaufort, Chukchi, and Bering seas (2011–2017). Ecology and Evolution, 10(12), 5595–5616.

Wolfinger, R. & O’connell, M. (1993). Generalized linear mixed models a pseudo-likelihood approach. Journal of Statistical Computation and Simulation, 48(3-4), 233–243.

Wood, S. N. (2017). Generalized additive models: an introduction with R. CRC press.

